# Visual Congruency Modulates Music Reward through Sensorimotor Integration

**DOI:** 10.1101/2024.07.30.605944

**Authors:** Lei Zhang, Yi Du, Robert J. Zatorre

## Abstract

There is emerging evidence that a performer’s body movements may enhance the music-induced pleasure of audiences. However, the neural mechanism underlying such modulation remains largely unexplored. This study utilized psychophysiological and electroencephalographic data collected from listeners as they watched and listened to manipulated vocal (Mandarin lyrics) and violin performances of Japanese and Chinese pop music. All participants were unfamiliar with the violin or Mandarin. The auditory and visual elements of the stimuli were either congruent (original recording) or incongruent (drawn from unrelated music videos). We found that congruent visual movements, as opposed to incongruent ones, increased both subjective pleasure ratings and skin conductance responses but only during vocal performances. Then, we examined the coherence between the music signal and sensorimotor Mu-band oscillatory neural activity and found that congruent visual movements enhanced Mu entrainment exclusively to vocal music signal. Further, mediation analysis demonstrated that neural entrainment to vocal music significantly mediated the visual modulation of music-induced pleasure. In conclusion, our study provides novel evidence on how congruent visual movements can heighten music-induced pleasure through enhanced sensorimotor integration.

## Introduction

Music consists of acoustic sequences that can give us pleasure (Zatorre, 2023). However, music often transcends purely auditory art; enjoyment frequently arises from both the auditory experience and the visual observation of performers’ gestures, whether live or via video. In addition to the audio part of music, performers’ body movements and facial expressions are also a part of their performance. While the audio aspects of music are well-recognized, the role of information conveyed visually in enhancing music-induced pleasure remains poorly understood.

Prior behavioral research has explored the impact of visual movement cues on music enjoyment. A meta-analysis of 15 studies quantified the average effect size of the enhancement in music performance appreciation under audio-visual conditions compared to audio-only conditions. It found a relatively consistent medium effect size increase in pleasure components was present (Platz and Kopiez, 2012). Moreover, several studies have suggested that visual cues may have a more significant influence on judgments about music performance than auditory cues (Griffiths and Reay, 2018; Tsay, 2013). These findings imply that the visual elements of music play a role in the experience of pleasure from music, at least at the behavioral level. However, further objective evidence is needed to substantiate these findings and to understand the potential physiological and neural mechanisms.

Skin conductance response (SCR), driven by sympathetic nervous system, measures phasic changes in the electrical conductivity of the skin (Braithwaite et al., 2013). SCR is often used as an objective indication of physiological arousal, and can be modulated by music-induced pleasure. When subjects hear music that they rate as pleasant, SCRs increase with increasing pleasurable states (Mas-Herrero et al., 2014; Salimpoor et al., 2009). However, no study has yet examined whether observing performers’ gestures could enhance the pleasure derived from music using objective physiological measures.

The enjoyment of music is also associated with movements in synchrony with the rhythm of the music. The shared affective motion experience (SAME) model proposes that music is perceived not only as an auditory signal but also through the intentional, hierarchically organized sequences of expressive motor acts behind the signal (Overy and Molnar-Szakacs, 2009). While listening to music, listeners may mimic the articulation or other movements of the performer, either overtly or covertly, facilitating the corepresentation and sharing of a musical experience between performer and listener (Dalla Bella et al., 2001; Gagnon and Peretz, 2003; Gomez and Danuser, 2007; Michalak et al., 2009; Overy and Molnar-Szakacs, 2009; Sievers et al., 2013). Neuroimaging studies have highlighted the engagement of the motor system during music listening, especially when the listener is proficient with the instrument being played (Gordon et al., 2018; Herholz et al., 2016; Lahav et al., 2007; Srinivasan et al., 2020), indicating that the motor system may play a role in aspects of perception.

Mu wave activity, a brain rhythm typically defined in the 8–13 Hz frequency band, provides a valid approach for studying the human motor system and sensorimotor integration with electroencephalogram (EEG) (Fox et al., 2016; Hobson and Bishop, 2017; Jenson et al., 2020). Reduced spectral power in the mu wave range, which originates in premotor areas and inferior parietal lobule, is observed during both action execution and action observation (Bonini et al., 2022; Fox et al., 2016, 2016; Hobson and Bishop, 2017; Jenson et al., 2020). Mu power suppression is also observed in music perception (Mercadié et al., 2014; Ross et al., 2022; Wang et al., 2023), and is further amplified when music perception is coupled with visual actions (Tanaka, 2021).

Recent studies using naturalistic continuous stimuli have begun investigating the neural tracking of continuous stimulus signals like speech envelope (Aller et al., 2022; Harding et al., 2019; Park et al., 2016). Cerebral-acoustic coherence (CACoh) analysis allows the measurement of the coherence between neural signals and continuous audio envelope signals (Harding et al., 2019; Peelle et al., 2013; Teng et al., 2024). Therefore, both Mu power suppression with naturalistic continuous stimuli and CACoh analysis can enable us to investigate how visual actions may modulate Mu entrainment to music amplitude envelope, and the correlation between neural entrainment to music and music-induced pleasure, which has never been done before.

In the present study, we recorded simultaneous electrodermal activity (EDA) and EEG signals when participants who did not know how to play violin listened to naturalistic vocal or violin music under four conditions: audio-visual congruent (AVc, where music and visual performance were matched), audio-visual incongruent (AVic, where audio tracks were switched across videos), visual only (VO), and audio only (AO) (Figure 1). We aimed to investigate whether visual congruency influences music-induced pleasure and whether this effect is mediated by sensorimotor mechanisms.

**Figure 1.**
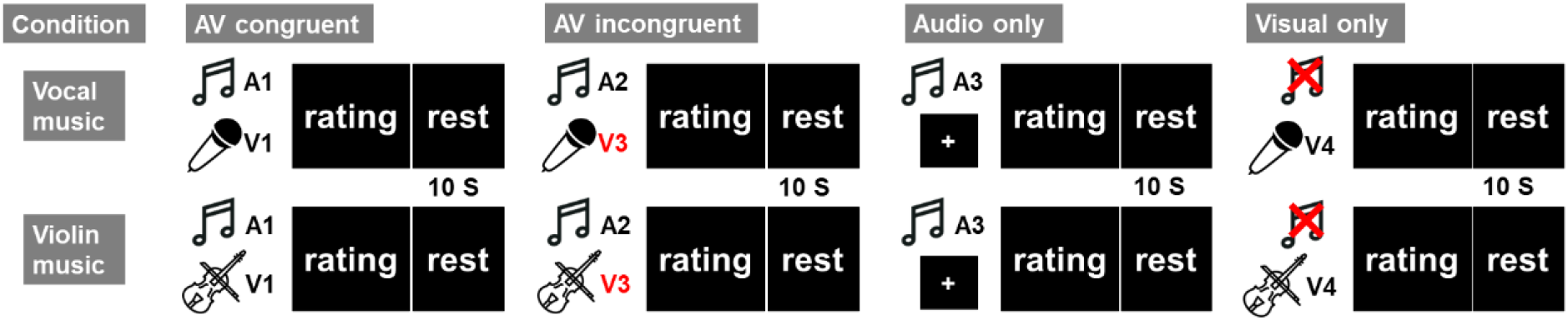
Experiment conditions. A denotes audio music stimuli and V denotes visual movements stimuli. AV congruent condition: Music stimuli and visual body movements stimuli were from the same video. AV incongruent condition: Music stimuli and visual body movements stimuli were from different video. Audio only condition: Only music stimuli was presented and a static cross was presented on the screen. Visual only condition: Only silent visual body movements movement video was presented.

We first analyzed subjective pleasant/liking ratings and EDA to assess the effect of visual congruency on musical pleasure using both subjective and objective indices. We then extracted Mu wave EEG activity (Mu power suppression and Mu entrainment to music signal) to investigate how visual congruency affects sensorimotor integration during music perception. Given our hypothesis that musical pleasure is modulated by visual congruency through sensorimotor integration, and considering that none of the participants could play the violin, we predicted that visual congruency would primarily influence the pleasure derived from vocal music, as participants could not utilize movement cues specific to playing the violin, such as fine finger movements but could interpret vocal movements, which are universally familiar. Lastly, we conducted correlation analyses and mediation analyses among visual congruency, musical pleasure, and Mu wave activity (power suppression and entrainment) to investigate their relationship during neural processing of music.

## Results

### Congruent visual information increases subjective pleasure ratings to music

After each music piece under AVc, AVic, and AO conditions, participants gave their liking, pleasure, arousal, and familiarity ratings to the music pieces. After each video under the VO condition, participants gave their liking ratings to the video.

One-way repeated measures analysis of variance (ANOVA) found that the main effects of visual conditions (AVc, AVic, and AO) on subjective pleasure ratings and liking ratings to music were significant only for vocal music (Pleasure: Vocal: *F*(2, 30) = 10.09, *p* < .001, 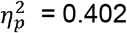; Violin: *F*(2, 30) = 1.19, *p* = 0.166, 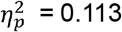; Liking: Vocal: F(2, 30) = 6.97, p = 0.003, 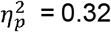; Violin: F(2, 30) = 2.23, p = 0.125, 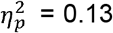). No significant main effect of visual conditions was found on arousal and familiarity ratings (Arousal: Vocal: *F*(2, 30) = 0.35, *p* = 0.705, 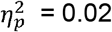 ; Violin: *F*(2, 30) = 1.93, *p* = 0.163, 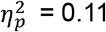; Familiarity: Vocal: F(2, 30) = 0.02, p = 0.985, 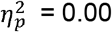; Violin: F(2, 30) = 0.71, p = 0.501, 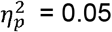). We also perform two-way repeated ANOVA (visual conditions × music type) to examine whether the effect of visual conditions on liking and pleasure ratings are different between vocal and violin music. However, no significant interaction effect was found (Pleasure: *F*(2, 60) = 0.88, *p* = 0.403, 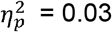 ; Liking: *F*(2, 60) = 0.83, *p* = 0.829, 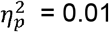).

As is shown in Figure 2a, post hoc analysis found that congruent visual information increased listeners’ pleasure ratings and liking ratings to vocal music compared to AVic condition (pleasure rating: *t*(30) = 3.87, *p*_corrected_ < .001, Cohen’s *d* = 0.79; liking rating: t(30) = 3.53, *p*_corrected_ = 0.004, Cohen’s d = 0.79). And no significant effect was found for violin music (pleasure rating: *t*(30) = 1.93, *p*_corrected_ = 0.191, Cohen’s *d* = 0.28; liking rating: t(30) = 2.09, *p*_corrected_ = 0.137, Cohen’s d = 0.31). To examine whether congruent visual information increased music-induced pleasure and liking ratings or incongruent visual information decreased music-induced pleasure and liking ratings, we compared pleasure and liking ratings between audiovisual conditions and AO condition (Wilcoxon signed rank tests were performed if the data violated the assumption of normality. Z-scores denote the statistics for Wilcoxon signed rank tests, and r values denote the effect size). Pleasure rating under AVc condition to vocal music was higher than that under AO condition (*Z* = 2.82, *p*_corrected_ = 0.015, *r* = 0.51), while liking rating did not increase under AO condition (*t*(30) = 2.04, *p*_corrected_ = 0.151, Cohen’s *d* = 0.42). However, pleasure rating and liking rating were not significantly different between AVic and AO conditions (pleasure rating: *t*(30) = 1.27, *p*_corrected_ = 0.642, Cohen’s *d* = 0.64; liking rating: AO: *Z* = 2.00, *p*_corrected_ = 0.138, *r* = 0.36). The results suggest that the impact of visual congruency on pleasure ratings to music can be attributed to the enhancement of these ratings by congruent visual information.

**Figure 2.**
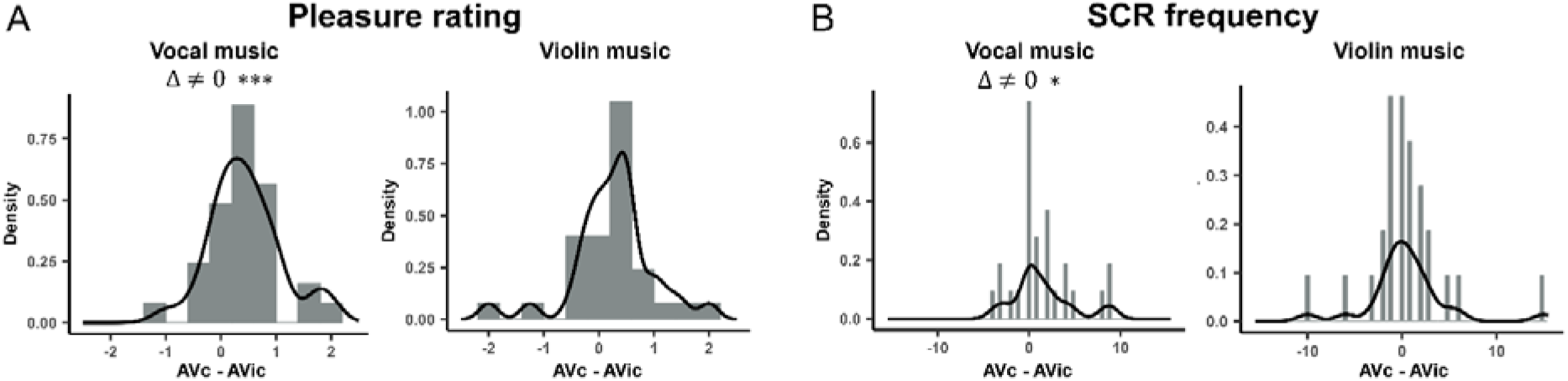
Histogram and density plot of the difference of subjective pleasure ratings and SCR frequency. Subjective pleasure ratings (A) and relative SCR frequency (B) were higher under AVc condition than AVic condition for vocal music but not for violin music.

### Congruent visual information increases skin conductance responses during music listening

SCR frequency and amplitude were extracted from EDA data to investigate the effect of visual congruency on electrodermal activity. As shown in Figure 2b, paired Wilcoxon signed rank test revealed that subjects showed significantly more SCRs under AVc condition than AVic condition when listening to vocal music (*Z* = 1.98, *p* = 0.048, *r* = 0.38), while no significant difference was found in violin music (Figure 2b). Subjects also showed marginally larger summed SCR amplitude when listening to vocal music with congruent visual movement compared to AVic condition (*Z* = 1.87, *p* = 0.061, *r* = 0.36), while no effect was found in violin music. No significant effect of music type on SCR difference between AVc and AVic conditions was found (SCR frequency: *Z* = 1.28, *p* = 0.201, *r* = 0.25; summed SCR amplitude: *Z* = 1.40, *p* = 0.162, *r* = 0.27).

### Visual information inhibits Mu-band power independently of congruency

Mu-band power was extracted for each visual condition to investigate whether the presence of visual information inhibited Mu-band power and whether visual congruency modulated Mu-band power during music perception. As is shown in Figure 3a, we found that Mu-band power under conditions with visual information was significantly inhibited compared to AO condition at almost all frontal, temporal, and occipital electrodes (AVc vs. AO, AVic vs. AO, *p*_fwe_ < 0.05) for both vocal and violin music. However, we did not find any electrodes that showed a significant difference in Mu power between AVc and AVic conditions.

**Figure 3.**
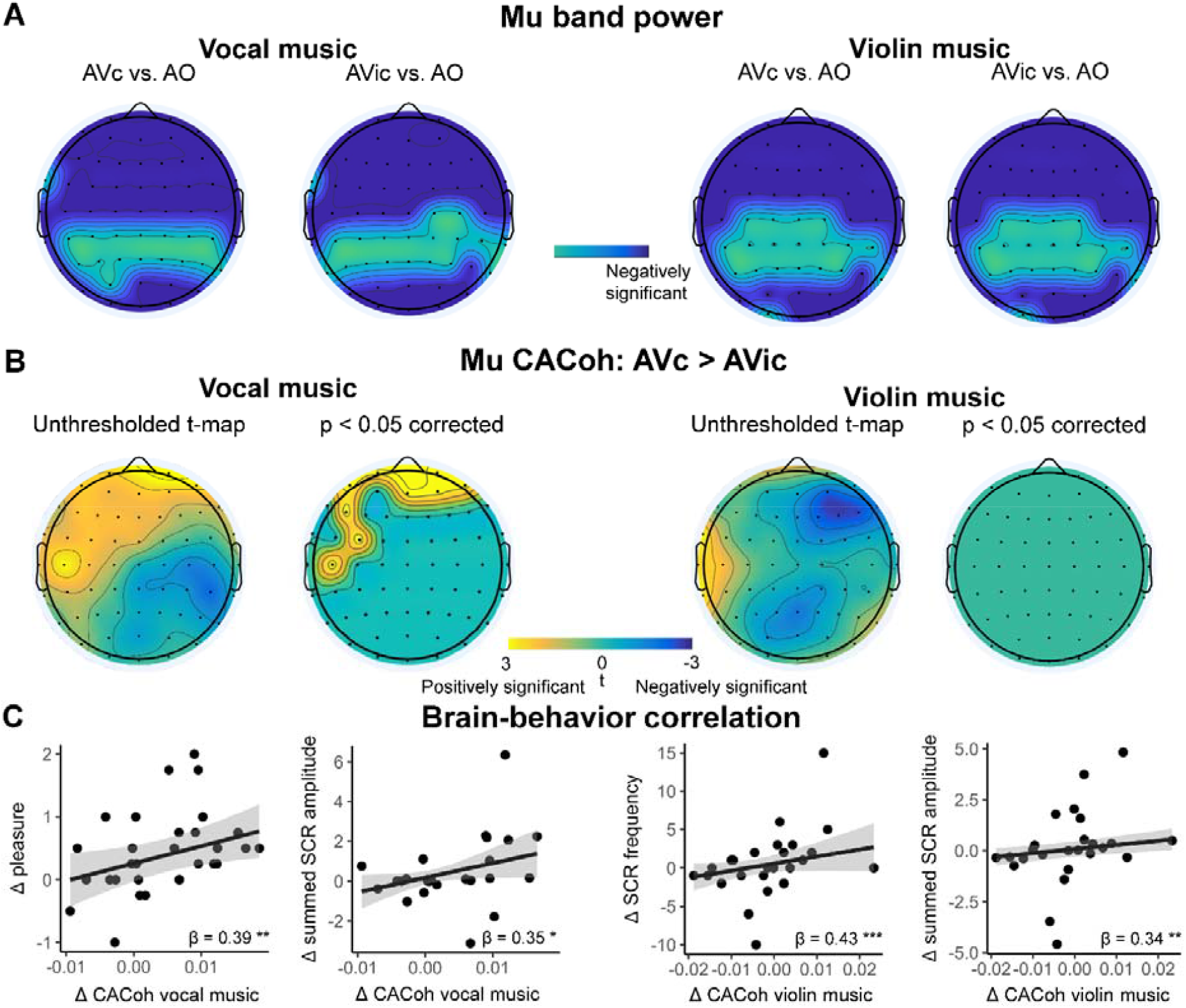
(A) Mu-band power was suppressed under AVc and AVic conditions compared to AO condition for both vocal and violin music. (B) Mu entrainment to music, as calculated by cerebral-acoustic coherence (CACoh) between Mu-band signal and music amplitude envelope, was enhanced under AVc condition compared to AVic condition only for vocal music. (C) Enhanced neural entrainment to music was correlated to greater subjective pleasure ratings and SCR indices for both vocal and violin music.

Additionally, Mu-band power was significantly suppressed at all central and frontal electrodes under AVc condition compared to AO condition during vocal music perception. However, for violin music, Mu-band power under AVc condition did not show significant suppression at Cz, C1, C2, and C3 electrodes, which typically showed Mu suppression when a person performed or observed an action (Hobson and Bishop, 2017), possibly because our participants do not know how to play the violin.

### Visual congruency modulates neural entrainment to vocal music in the Mu band

In addition to Mu-band power, CACoh analysis was performed to investigate the effect of visual congruency on the neural entrainment to the music signal in the Mu band. As shown in Figure 3b, we found that CACoh vocal music was significantly greater at frontal electrodes (FP1, FP2, AF7, AFz, AF4, AF8, F5, FC3, and C5) under AVc condition than AVic condition (*p*_fwe_ < 0.05). However, no electrodes showed a significant effect of violin music CACoh.

Robust linear regression analysis was conducted to investigate the association of the music CACoh difference at electrodes that showed a significant visual congruency effect with the SCR difference, and with subjective rating difference between AVc and AVic conditions. We found that the average increased CACoh across significant channels could significantly predict the increased pleasure ratings in vocal music (Figure 3c, β = 0.39, p = 0.001). Also, increased CACoh could predict larger summed SCR amplitude for vocal music (β = 0.35, p = 0.034).

### Greater Mu entrainment to violin music with congruent visual information is correlated to increased music-induced pleasure and Mu suppression

We did not find a visual congruency modulation effect to violin music-induced pleasure and neural entrainment generally. However, considering individual differences in audiovisual violin music perception, we hypothesized that subjects who showed greater Mu suppression or Mu entrainment to music under visual congruent condition compared to visual incongruent condition could demonstrate greater music-induced pleasure. Therefore, we performed robust linear regression analysis to examine whether Mu power difference across electrodes that showed significant suppression in audiovisual vocal music but not in audiovisual violin music (Cz, C1, C2, and C3) or average CACoh difference at electrodes that showed greater neural entrainment to vocal music under AVc condition (FP1, FP2, AF7, AFz, AF4, AF8, F5, FC3, and C5) could predict violin music-induced pleasure difference. And we also examine the relationship between the average CACoh difference and Mu power difference.

Robust linear regression results showed that greater Mu entrainment under AVc condition compared to AVic condition predicted greater summed SCR amplitude, SCR frequency, and Mu suppression for violin music (Figure 3c, summed SCR amplitude: β = 0.34, p = 0.005; SCR frequency: β = 0.43, p < 0.001; Mu suppression: β = -0.36, p < 0.001).

Additionally, greater Mu suppression was correlated with increased subjective arousing ratings (β = -0.16, p = 0.021), and longer musical training years could marginally predict greater Mu suppression (β = -0.15, p = 0.065).

### Increased neural entrainment to vocal music mediates increased music-induced pleasure rating with congruent visual information

Since we found that visual congruency modulated both Mu entrainment to music and music-induced pleasure to vocal music, we explored whether enhanced entrainment to music mediates increased music-induced pleasure under AVc condition. The electrodes that showed significantly greater Mu entrainment to vocal music were included in the within-subject mediation analysis. As shown in Figure 4a, we found a significant indirect effect of visual congruency on subjective pleasure ratings through increased mean music CACoh value across all significant electrodes (Figure 4a, visual congruency → average CACoh value → subjective pleasure rating, 95% CI, 0.01 to 0.23). Additionally, we found a full mediation effect of the mean Mu entrainment, since visual congruency was not correlated with pleasure rating after controlling the mean Mu entrainment (from β = 0.27, p < 0.001 to β = 0.17, p > 0.05). These results indicated that the Mu entrainment fully explained the visual modulation on subjective pleasure rating.

**Figure 4.**
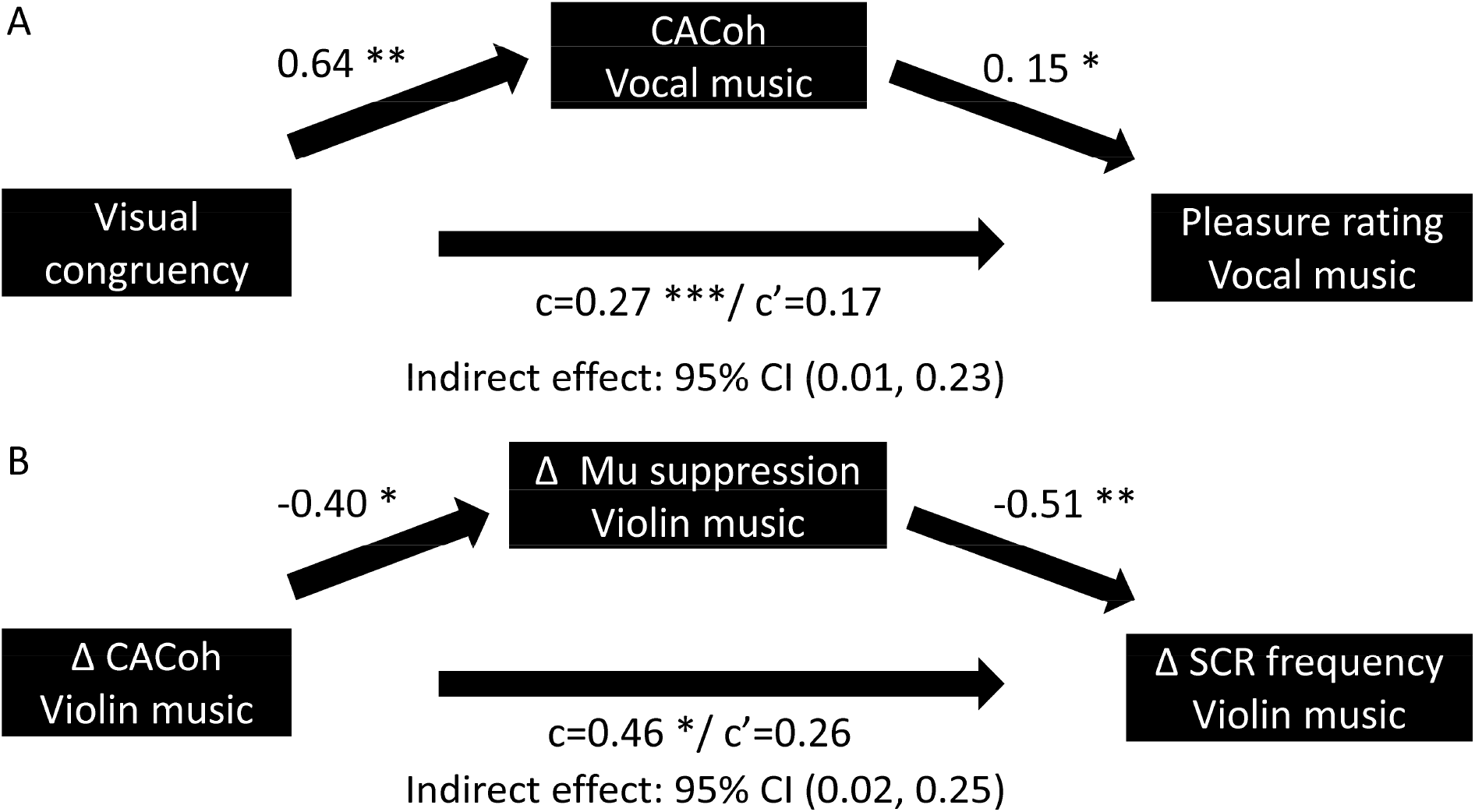
Within-subject mediation analysis results. (A) Mu entrainment to vocal music mediated the visual modulation on subjective pleasure ratings. (B) Enhanced Mu power suppression to violin music under AVc condition was the significant mediator of the correlation between enhanced Mu entrainment and greater SCR frequency to violin music under AVc condition compared to AVic condition. CACoh: Cerebral-acoustic coherence.

### Increased Mu suppression mediates the association between increased neural entrainment to music and music-induced SCR change

In violin music, we discovered a significant correlation between the difference in Mu entrainment, SCR frequency, and the summed SCR amplitude difference under AVc and AVic conditions, as well as the difference in Mu suppression. We also conducted the mediation analysis to explore further relationships between neural and psychophysiological indices. As shown in Figure 4b, we found a significant full mediation effect of Mu suppression between neural entrainment to music and SCR frequency difference (Δ CACoh value → Δ Mu suppression → Δ SCR frequency, 95% CI, 0.02 to 0.25, from β = 0.46, p = 0.019 to β = 0.17, p > 0.05). These results suggest that increased Mu suppression fully explains the relationship between neural entrainment to music and the physiological difference. However, the mediation effect was not significant when neural entrainment was set as the mediator (Δ Mu suppression→ Δ CACoh value → Δ SCR frequency, 95% CI, -0.26 to 0.05). Also, we did not find a significant mediation effect when summed SCR amplitude difference was set as the dependent variable (Δ CACoh value → Δ Mu suppression → Δ summed SCR amplitude, 95% CI, -0.03 to 0.40; Δ Mu suppression→ Δ CACoh value → Δ summed SCR amplitude, 95% CI, -0.30 to 0.10).

## Discussion

In this study, we present evidence that congruent performer movements enhanced listeners’ musical rewards compared to incongruent movements, at both behavioral and physiological levels using naturalistic music pieces. Such enhancement is likely attributable to the facilitation of sensorimotor integration among listeners, as indicated by Mu entrainment to music during the listening experience. Notably, all participants were non-violin players, and these findings were exclusively observed in vocal music but not violin music. This further substantiates that enhanced sensorimotor integration serves as a crucial neural mechanism for augmenting musical rewards through visual movement information.

Several previous behavioral studies have found that music performance appreciation, as subjectively rated by listeners, increases when observers view congruent performers’ movement (Vines et al., 2011, 2006; Tsay, 2013; Eaves et al., 2020; Griffiths and Reay, 2018; Platz and Kopiez, 2012). Consistent with this phenomenon, we found that listeners’ subjective pleasure ratings were modulated by the congruency of visual information, with increased pleasure under congruent conditions. Furthermore, SCRs, which serve as reliable objective indicators of arousal and/or reward levels in response to music stimuli (Martínez-Molina et al., 2016; Mas-Herrero et al., 2014; Salimpoor et al., 2009; Solberg and Dibben, 2019; Zatorre, 2015), were found to increase in frequency and amplitude under congruent visual conditions. These results support the hypothesis that visual congruency between the performer’s action and the music significantly heightens musical pleasure.

Previous studies have also found improved speech perception when observers simultaneously view congruent lip movements (Park et al., 2016; Zhang and Du, 2022; Zion Golumbic et al., 2013), which provides both content information and predictive timing of the speech (Peelle and Sommers, 2015). Several MEG and EEG studies have found better neural tracking of speech amplitude envelope when listening to speech with congruent lip movements (Crosse et al., 2016, 2015; Giordano et al., 2017; Park et al., 2016; Zion Golumbic et al., 2013). Analogous effects were observed in the present study regarding music envelope tracking through coherence analysis, suggesting that similar mechanisms, that is, performers’ movements, may provide content information and predictive timing of the vocal music, may enhance neural entrainment to music.

The role of the motor system in music-induced pleasure remains underexplored. A recent meta-analysis failed to find a significant association between cortical motor areas and music-induced pleasure (Mas-Herrero et al., 2021). Previous fMRI studies have shown that the motor system is involved in rhythmic information processing (Gordon et al., 2018), and several studies have found correlations between cortical and subcortical motor areas and beat-related pleasure (Kornysheva et al., 2010; Matthews et al., 2020; Trost et al., 2014). In vocal music perception, visual vocal motor information provides music features like timing, pitch, and amplitude. Listeners could employ these visual cues to better predict the upcoming music, thereby increasing predictability and reducing uncertainty with congruent visual movements of the performers. Previous studies investigating how music surprisal and entropy interact with pleasure rating found that music with intermediate levels of complexity (surprisal) and uncertainty (entropy) received higher pleasure ratings (Cheung et al., 2019; Gold et al., 2019). Therefore, we speculate that clearer content and greater predictive information perceived through congruent visual information leads to a better internal representation and, consequently, greater pleasure for the listeners.

In addition, greater coherence between neural activity and the music envelope in the Mu band was only observed in vocal music, not in violin music, over sensorimotor and frontal electrodes. None of the participants in this study could play the violin, while all could sing to some extent. Mu-band EEG signals reflect activities of the human mirror neuron system (Hobson and Bishop, 2017). Recent neuroimaging and neural modulation studies have found that the recruitment of the motor system is important for speech perception, especially under adverse conditions (Du et al., 2014; Hickok et al., 2011; Liang et al., 2023; Murakami et al., 2015; Zhang et al., 2023; Zhang and Du, 2022). Although not crucial, previous studies have found that motor system is also involved during music listening, especially when people have acquired knowledge of playing the musical instrument (Gordon et al., 2018; Herholz et al., 2016; Lahav et al., 2007; Srinivasan et al., 2020). Several musical training experiments have found that after training, listeners showed greater sensorimotor integration during music listening (Herholz et al., 2016; Herholz and Zatorre, 2012; Srinivasan et al., 2020; Wollman et al., 2018; Zatorre et al., 2007). These findings may explain why listeners benefited from vocal music production video. With knowledge of vocal music production, they can extract more information from congruent visual vocal movements. Conversely, since the participants did not know how to play the violin, they could not derive as much music information from specific violin-playing motions, rendering visual congruency less influential on violin music perception.

Although we did not find a significant modulation of visual congruency on violin music processing or pleasure, some subjects may still gather information from violin playing movements, such as bow movements that are correlated with note onset and sound volume. Robust regression analysis revealed a significant correlation between enhanced Mu entrainment and violin music-induced pleasure, indicating that listeners who exhibited greater Mu entrainment enhancement to violin music were more likely to experience greater violin music-induced pleasure with congruent visual movements. These results imply that sensorimotor integration plays an important role in visual enhancement of music-induced pleasure.

Mu band power is typically suppressed when observing or imitating actions of others (Hobson and Bishop, 2017). In this study, we found Mu suppression under both AVc and AVic conditions compared to the AO condition, with no difference between AVc and AVic conditions. Instead, the finding that visual congruency significantly modulated the coherence between Mu band neural activities and music signals provides a new approach to investigate the role of Mu-band brainwave in sensory perception.

There are several limitations in this study. First, due to the use of natural stimuli, the effect of performers’ general movements, such as body sway and fine movements in violin playing, cannot be separated. To address this issue, future studies could employ motion capture to record audiovisual music stimuli, allowing separate investigation of different motion components. In addition, the relatively poor spatial resolution of EEG signals prevents us from identifying specific brain areas in the frontal lobe that contribute to the enhanced brain entrainment to music and music-induced pleasure. In future studies, neuroimaging techniques with higher spatial resolution like MEG and fMRI could be employed to obtain more accurate spatial information.

In sum, this study provides empirical evidence that music-induced pleasure can be modulated by observing performers’ body movements, as reflected in both subjective ratings and skin conductance responses. Since performers’ movements provide information about music production, we propose that sensorimotor integration of the audience is enhanced during music perception, leading to better predictions and reduced uncertainty, factors that are believed to influence pleasure (Zatorre, 2023). Enhanced sensorimotor integration is a significant factor through which an audience experiences greater music-induced pleasure with congruent visual movements compared to incongruent visual movements.

## Method

### Participants

The sample size was estimated using power analyses (α = 0.05, power = 0.8) in G*Power 3.1 (Faul et al., 2009). Using a two-tailed paired t-test, 33 participants would be sufficient to detect the effect size (Cohen’s d = 0.51) estimated by a previous meta-analysis (Platz and Kopiez, 2012) with 80% power. Based on the power analysis and experimental design, 32 participants (18 females) were recruited for this study. One subject was removed from further analysis due to fatigue. Additionally, the first two subjects’ EEG data were invalid due to a technical error and were discarded from further analysis, although their behavioral and EDA data were retained. Four subjects’ EDA data were excluded from the analysis because they exhibited no skin conductance responses after taking a deep breath and holding it for several seconds, indicating non-responsiveness in terms of EDA (Braithwaite et al., 2013).

All the participants (18 females, 18 – 33 years old) were right-handed, non-Mandarin speakers, without any neurological or psychiatric disorders, and had no more than 2 years of training in string instruments or vocals (although some had training in other instruments). The Barcelona Musical Reward Questionnaire (BMRQ) scores of all participants were above 65 (indicating no music anhedonia) (Mas-Herrero et al., 2013). All participants had signed the written consent approved from the Institutional Review Board of McGill University.

### Stimuli and procedure

Twenty naturalistic music pieces (10 vocal and 10 violin) were selected from YouTube and Bilibili. These music pieces were all Japanese or Chinese pop music, with vocal music lyrics in Mandarin. The pieces were selected so that they would likely be unfamiliar to our Western listeners who did not speak Mandarin. Consequently, any observed effects of congruency can be attributed solely to lower-level cues, such as the movements of the mouth, fingers and arms, and their synchronization with the musical sounds. The videos only contained the performer’s movements against a static background, with no other musical accompaniment. Dynamic parts of the videos, such as rolling titles, were blurred by a Gaussian function to ensure they were invisible to the subjects. Music videos were trimmed to approximately 80-second segments. A pilot study was conducted to further refine the selection of music pieces, where 13 participants were asked to rate their liking of the music. Eight pieces were selected based on the liking ratings and the length of the music, and were used in the psychophysics and EEG experiment. The average sound intensity of each audio was normalized according to the root mean square value, and the sampling rate was 48 kHz. The video resolution was 1280 × 720 pixels, with a frame rate of 30 frame/s. The video and audio parts of the clips were combined according to 4 conditions (see Figure 1 for the experimental design).

#### Audiovisual congruent condition

(AVc). Music and visual performance were matched (i.e. the intact video was used);

#### Audiovisual incongruent condition

(AVic). Music and visual performance were mismatched by switching audio tracks across videos (see below);

#### Audio-only condition

(AO). Only music was played with a static cross at the center of the screen;

#### Visual-only condition

(VO); Only video was played without music.

Note that in the AVic condition, vocal video was combined only with vocal music, and violin video only with violin music. The same videos and music from the AVc condition were used to generate the AVic stimuli. Each pair of pieces was always combined with each other. For example, if vocal music 1 was combined with vocal video 2, vocal music 2 was combined with vocal video 1, and vice versa. Pairs of music pieces with similar durations were chosen to be combined with each other. If one peice was longer or shorter, the video was trimmed to match the music duration. Each trial began with a 3-second static image, which was the first frame of the video.

The experiment consisted of 8 blocks (violin/vocal × 4 conditions). Each vocal or violin block contained 4 trials with different conditions. After each trial, participants rated (0-9) their liking, pleasure, arousal, and familiarity with the music if music was presented, and their liking (0-9) of the visual performance if the visual performance was presented. Conditions were counterbalanced using a Latin square design. Music type was also balanced using an ABBA design (A denoted a violin-vocal block sequence, and B denoted a vocal-violin block sequence). Therefore, an ABBA sequence denoted violin/vocal/vocal/violin/vocal/violin/violin/vocal block). ABBA sequences were presented to half of the subjects, and BAAB sequences were presented to the other half.

### Behavioral data analysis

Behavioral responses to vocal and violin music were analyzed separately. We performed a one-way repeated measures ANOVA to investigate the effect of visual conditions on ratings for vocal and violin music. Greenhouse–Geisser correction was performed if the sphericity assumption was violated. Post hoc analysis further examined the difference between specific conditions (AVc vs AO, AVic vs AO, AVc vs AVic). Wilcoxon signed rank tests were performed if the data violated the assumption of normality, as assessed by the Shapiro-Wilks normality test. Bonferroni correction was applied for multiple comparisons. Statistical analysis was conducted in R with the package bruceR (Bao, 2020) and visualized with ggplot2 (Wickham, 2009).

### EDA data acquisition and analysis

EDA data were collected using the BIOPAC MP150 system with the EDA 100C amplifier module. Two electrodes with isotonic gel were placed on the volar surfaces of the distal phalanges of the right index and middle finger. The sampling rate was 2000 Hz, and the gain was set to 5 μS/V.

To keep the consistency of the data length of each condition, the first 77 seconds of each trial were analyzed. EDA was analyzed using Neurokit2 (Makowski et al., 2021), a python toolbox, and custom scripts. Default parameters were used. The number and summed amplitude of SCR in each condition were extracted. Note that, only SCRs that reached a peak between 2 seconds after the onset and the end of the music were considered, as the latency of SCR is about 2 seconds.

To account for baseline differences in SCR activity among subjects, the AO condition was treated as the baseline, and the relative difference in SCR indexes under AVc and AVic conditions entered the latter statistical analyses.

Two-sampled paired Wilcoxon signed rank tests were performed to compare SCR indexes between AVc and AVic conditions for vocal and violin music separately, as the differences violated the normal distribution assumption (Shapiro-Wilk normality test, both p < 0.042).

### EEG data acquisition and processing

EEG data were acquired from 64 Ag/AgCl electrodes using Brain Products actiCAP active electrode system, including 2 electrooculogram (EOG) electrodes. Impedance of all electrodes was kept below 25 kΩ, sufficient for high-quality data collection according to the actiCAP system manual. All channels were referenced to the average of the left and right mastoid channels (M1, M2).

EEG data analysis was conducted using the Fieldtrip toolbox (Oostenveld et al., 2011) and custom scripts. Similar to EDA data analysis, only the first 77 seconds of each trial were analyzed. Raw EEG data were first bandpass filtered from 1 to 40 Hz, and downsampled to 90 Hz. Independent component analysis (ICA) was performed to remove the artifacts from eye blinks, eye movements, and heartbeat. Mastoid channels and EOG channels were removed in the latter analysis.

### Spectral analysis

To investigate how visual performance affects Mu wave activity, spectral analysis was performed at each electrode. Preprocessed data were segmented into non-overlapping 1-second intervals. Power spectrum was computed and averaged across each condition using the function “ft_freqanalysis” provided in FieldTrip. Frequencies of interest were from 1 to 40 Hz in steps of 0.5 Hz. Spectral power from 8 to 13 Hz was averaged to get the overall mu wave power under each condition.

### Cerebral-acoustic coherence (CACoh) analysis

CACoh analysis calculated cross-spectrum coherence between neural signals and music amplitude envelopes, which allows us to quantify phase-locked neural responses to acoustic signals at specific frequencies (Harding et al., 2019; Peelle et al., 2013; Teng et al., 2024). Audio signals were filtered using a bank of fourth-order gammatone filters with 16 center frequencies logarithmically spanning from 100 Hz to 4000 Hz. Hilbert transformation extracted the amplitude envelope of each band. The average of the 16 amplitude envelopes was downsampled to 90 Hz and bandpass filtered from 1 to 40 Hz to match the EEG signal for CACoh analysis.

We employed the function ‘mscohere’ in MATLAB to calculate magnitude-squared coherence. Magnitude-squared coherence between EEG signals at each electrode and the amplitude envelope of the music was estimated from 1 to 40 Hz in steps of 0.5 Hz across each condition. Coherence estimate was averaged between 8 to 13 Hz to assess the neural entrainment of Mu waves to the music.

### Cluster-based permutation test

The cluster-based permutation test in FielfTrip was performed to correct for multiple comparisons in spectral and CACoh analyses. Two-sided paired t-tests assessed differences in CACoh values between AVc and AVic conditions at each electrode. The sum of the T-statistics for statistically significant electrodes with a cluster was calculated as the cluster-level statistic. 5000 Monte Carlo-based permutations were conducted to obtain the permutation distribution. The cluster-level alpha was set to 0.05.

### Regression and mediation analysis

Robust linear regression analysis was performed to investigate the correlation between neural indices and music-induced pleasure using the “robustbase” package in R (Maechler et al., 2023).

Within-participant mediation analysis was performed using the “JSmediation” package in R (Yzerbyt et al., 2018) to investigate whether music processing mediated the enhanced pleasure with valid visual movements. For vocal music, the average CACoh difference of significant electrodes between AVc and AVic conditions was set as the mediator variable. Visual congruency was the binary predictor, and pleasure rating was the dependent variable. For violin music, the average CACoh difference of the electrodes were the predictor, SCR frequency or summed amplitude difference was the dependent variable, and Mu suppression difference was the mediator. Note that, the “JSmediation” package reported original regression coefficients instead of robust regression coefficients for mediation analysis. Confidence intervals of the within-participant indirect effect were computed using the Monte Carlo method with 5000 iterations. The alpha threshold was set to 0.05.

## Acknowledgments

We thank Alberto Ara, Arielle Rabinowitz, Kazuma Mori, and Oscar Bedford for help with experiment setup.

## Author contributions

Conceptualization: L.Z., and R.J.Z. Methodology: L.Z., and R.J.Z. Analysis: L.Z. Investigation: L.Z. Writing—original draft: L.Z. Writing—review and editing: L.Z., Y.D., and R.J.Z. Visualization: L.Z., and R.J.Z. Supervision: Y.D., and R.J.Z.

## Data and materials availability

All data needed to evaluate the conclusions in the paper are present in the paper. The behavioral, EDA, and EEG data that support the findings of this study are available on OSF.

